# GSK3B inhibition partially reverses brain ethanol-induced transcriptomic changes in C57BL/6J mice: Expression network co-analysis with human genome-wide association studies

**DOI:** 10.1101/2025.04.03.647116

**Authors:** Sam Gottlieb, Dustin Zeliff, Brennen O’Rourke, Walker D. Rogers, Michael F. Miles

**Affiliations:** Department of Pharmacology and Toxicology; Program in Neuroscience; Department of Human and Molecular Genetics; VCU Alcohol Research Center Virginia Commonwealth University, Richmond VA 23298-USA

## Abstract

Alcohol use disorder (AUD) is a chronic behavioral disease with greater than 50% of its risk due to complex genetic contributions. Existing pharmacological and behavioral treatments for AUD are minimally effective and underutilized. Animal model behavioral genetics and human genome-wide association studies have begun to identify individual genes contributing to the progressive compulsive consumption of ethanol that occurs with AUD, promising possible new therapeutic targets. Our laboratory has previously identified *Gsk3b* as a central member in a network of ethanol-responsive genes in mouse prefrontal cortex, which altered ethanol consumption with genetic manipulation and was also significantly associated with risk for alcohol dependence in human genome-wide association studies. Here we perform detailed brain RNA sequencing transcriptomic studies to characterize a highly specific and clinically available GSK3B pharmacological inhibitor, tideglusib, as a possible therapeutic for clinical trials on treatment of AUD. A model of chronic intermittent ethanol consumption was used to study gene expression changes in prefrontal cortex and nucleus accumbens in the presence or absence of tideglusib treatment. Multivariate analysis of differentially expressed genes showed that tideglusib largely reversed ethanol-induced expression changes for two prominent clusters of genes in both prefrontal cortex and nucleus accumbens. Bioinformatic analysis showed these genes to have prominent roles in neuronal functioning and synaptic activity. Additionally, mouse brain differential gene expression data was analyzed together with human protein-protein interaction and genome-wide association studies on AUD to derive networks responding to tideglusib and relevant to human genetic risk for alcohol dependence. These studies identified discrete networks significantly enriched with genes provisionally associated with AUD, and provide key information on central hubs of such networks. Together these studies document tideglusib as a major modulator of chronic ethanol consumption-evoked brain gene expression signatures, and identify possible new targets for therapeutic modulation of AUD.

## Introduction

The risk to develop an AUD is estimated to be 50% attributable to genetic predisposition[1]. For example, deleterious single nucleotide polymorphisms (SNPs) in the genes encoding for ethanol metabolism, alcohol dehydrogenase (ADH) and aldehyde dehydrogenase (ALDH), alter the risk of AUD[2]. Human genome-wide association studies (GWAS) have emerged as a growing tool in identifying additional SNP variants correlated with AUD, providing insight into potential novel therapeutic targets[3, 4]. The GWAS approach examines individual SNPs across the genome, agnostic to any functional relevance, and by comparing the frequency of these variants in affected versus unaffected populations, regions of the genome associated with phenotypic variation can be pinpointed. Most recently, problematic alcohol use GWAS have identified 110 independent risk variants in humans[5]. However, while GWAS is a powerful tool, it requires very large subject numbers to overcome the multiple test corrections which result from probing millions of different SNPs across the genome. Due to the polygenic nature of AUD pathophysiology, where many variants each contribute moderate effects in and of themselves, there is a possibility of critical genes in the ethanol regulatory pathway failing to meet the strict criteria for significance in GWAS.

To study the effects of many genes on the etiology of AUD, researchers have utilized genome-wide expression profiling techniques such as microarray or RNA sequencing (RNAseq)[6–10]. Such genome-wide studies initially aimed to identify a small, targeted list of genes which were responsible for the heritability or mechanisms of AUD, however these methods soon revealed large, complex lists of genes from human, animal model or in vitro analyses. To reduce the complexity of such results, investigators often utilize data reduction techniques such as clustering or expression network analyses such as <u>w</u>eighted <u>g</u>ene correlation <u>n</u>etwork <u>a</u>nalysis (WGCNA). Such networks can then more productively be analyzed for functional over-representation, giving insight into important signaling pathways. Furthermore, within expression networks, “hub” genes are central members which display high interconnectivity to the other network members through expression correlates. Therefore, manipulation of a hub gene is predicted to strongly influence expression changes within its connected gene network. One such hub gene identified in an alcohol-related network using this approach was *Gsk3b*[6, 11], which was identified as a central member in a network of ethanol-responsive genes identified in mouse PFC and was also significantly associated with risk for alcohol dependence in human GWAS data[12].

Bioinformatic analysis of gene expression and AUD mechanisms can be expanded further by utilizing approaches that directly integrate different types of biological or genetic information, oftentimes across different species. An example of such an approach is Edge-weighted Dense module search for genome-wide association studies and gene expression profiles (EW-dmGWAS)[13]. EW-dmGWAS allows for identification of AUD-associated networks by using (for example) a combination of protein-protein interaction data, human GWAS, and mouse ethanol gene expression data. These results can then be scored using an overrepresentation analysis to determine the strength of their correlation to genes with a known association to AUD, providing novel insight for mechanisms that contribute to alcohol dependence[14].

Our prior studies in mouse models identified GSK3β as a potential target for AUD treatment, and more recent work has shown that a clinically available GSK3β inhibitor, tideglusib, potently decreases ethanol intake during both binge drinking and prolonged consumption. However, the mechanism through which GSK3β regulates ethanol consumption is unknown and warrants study to identify additional potential targets for intervention in AUD.

Here we perform a detailed RNA sequencing analysis (RNA-seq) in both prefrontal cortex (PFC) and nucleus accumbens (NAc) of C57BL/6J mice with a history of prolonged ethanol versus water only consumption in the presence or absence of tideglusib. We hypothesized GSK3β inhibition via tideglusib would ameliorate ethanol-induced changes in differential gene expression and would thus both identify the extent to which GSK3β signaling impacts ethanol-responsive gene expression, and verify the molecular actions of tideglusib by functional analysis of gene expression patterns. Additionally, we superimposed this RNA-seq data on human GWAS and human protein-protein interaction data using EW-dmGWAS to identify gene networks most relevant to risk for AUD. Our studies thus identify gene networks highly regulated by ethanol consumption that are also modulated by GSK3β inhibition and contain genes conveying risk for AUD in humans. This work provides further insight into mechanisms underlying AUD and how GSK3β inhibition could act as a therapeutic in AUD treatment.

## Materials and Methods

### Ethics Statement

Rodent animal studies and procedures were approved by the Institutional Animal Care and Use Committee of Virginia Commonwealth University and followed the NIH Guide for the Care and Use of Laboratory Animals.

### Animals

Adult male C57BL/6J mice were purchased from Jackson Laboratories (Bar Harbor, ME). Mice were housed in a temperature and humidity-controlled room in accordance with the Association for Assessment and Accreditation of Laboratory Animal Care approved animal care facility and had ad libitum access to food and water. Animals were housed on utilized Tekland Laboratory Grade Sani-Chips bedding and kept on a reverse 12hr light cycle. All animals in this study were used as a part of Experiment 2 in a recent preprint publication [15] and manuscript under review (Gottlieb et al. 2025).

### Drugs

Ethanol (EtOH) for drinking studies was prepared as a 20% (v/v) solution in tap water and provided to animals for voluntary oral consumption. The GSK3β inhibitor tideglusib (Selleck Chemicals, Houston, TX) was prepared for gavage by suspension at 20 mg/ml in corn oil by vortexing followed by bath sonication at 40° C for 60 minutes. Two separate gavage treatments of 100 mg/kg tideglusib were administered four hours apart on drinking days.

### Intermittent ethanol access, two-bottle choice (IEA)

Male mice underwent IEA for nine weeks as recently described [15]. Animals were habituated to IEA for three weeks. Mice were then acclimated to gavage treatments with two days saccharin administration followed by two days vehicle habituation before beginning gavage of 100 mg/kg tideglusib or equivalent volume vehicle on week five, after which effects on consumption and ethanol preference were measured for four additional weeks. Two separate gavage treatments were administered each drinking day. The first treatment occurred two hours prior to ethanol access and the second occurred four hours following the first, occurring immediately after the two-hour binge read.

### RNA sequencing analysis

A subset of mice was selected randomly (n=4/group) and were euthanized 24 hours after their last ethanol access to compare gene expression between animals +/- EtOH and +/- tideglusib. PFC and NAc were collected and sent for sequencing at the VCU Genomics Core. Paired end counts were analyzed for differential expression analysis across any two treatment groups using DESeq2. Genes with median counts <1 across all samples were filtered out of the data and significantly differently expressed genes (DEGs) were determined using an uncorrected p<0.05 and a log-fold change (LFC) of LFC>0.1 or LFC<-0.1. To generate hierarchical clustering across all four treatment groups, transcript counts in each brain region were variance-stabilizing transformed with the vst() function from DESeq2 (v. 1.40.2). Genes which met DEG criteria between any two experimental groups were included (PFC n=1664; NAc n=1838). Gene-wise z-scores were calculated across all samples within each brain region separately, and based upon gene expression patterns, hierarchical clustering was used to generate five clusters. Results from comparison of ethanol-vehicle treated animals vs. ethanol-tideglusib treatment are discussed in detail in another publication (Gottlieb and Miles, manuscript in review). Functional over-representation analysis of DEGs was performed using ToppGene[16].

### GWAS

For the human GWAS dataset we used a composite AUD GWAS dataset created by combining UKBiobank[3], the Psychiatric Genome Consortium[17], and GSCAN[18] results provided by Dr. B. Todd Webb. Genes that had a GWAS association p-value ≤10E-6 with alcohol consumption phenotypes were considered nominally significant in performing over-representation analysis on EW-dmGWAS networks.

### Edge-weighted, dense module GWAS (EW-dmGWAS)

The R package EW-dmGWAS[13] (v.3.0) was used to identify small, constituent networks (edge weighted modules) within the background network of binary protein-protein interactions (PPI). The PPI network was derived from PINA[19] (v.2.0), and the *Homo sapiens* set was used as it has 166,776 binary interactions compared to 13,865 for *Mus musculus*[19, 20]. Protein symbols were matched to the corresponding gene symbols using Uniprot identifiers[21]. RNA seq feature counts from the ethanol/vehicle (EV), water/vehicle (HV), ethanol/tideglusib (ET), and water/tideglusib (HT) treated mouse mPFC were used for the edge-weights.

For the purposes of this chapter, “network” will refer to the background PPI framework and “module” will refer to the resulting groups of genes identified by the EW-dmGWAS analysis. Higher node-weights represent more significant GWAS p-values, and higher edge-weights represent a larger difference in the correlation of the expression of two respective genes (nodes) between two different treatment groups (e.g., EV versus ET). This is calculated by taking the difference of correlations in expression values of the two genes. The module score algorithm incorporated edge- and node-weights, which were each weighted to prevent bias towards representation of nodes or edges in module score calculations. Such bias could cause some modules to be identified based almost solely on edge-weights or node-weights, as opposed to the two combined, which would defeat the purpose of integration. The respective weighting depends upon a parameter (λ) which is calculated prior to module searching, based on the entire set of node- and edge-weights and used across all module score calculations, as part of the EW-dmGWAS algorithm.

Higher module scores represent higher edge- and node-weights. Genes were kept in a module if they increased the standardized module score (Sn) by 0.5%. Sn corresponding to a permutation-based, empirical quantile false discovery rate (qFDR)<0.05 were considered significant. A significant Sn (i.e., more significant qFDR values) indicates a module’s constituent genes are more highly associated with alcohol consumption in humans, and their interactions with each other are more strongly perturbed by chronic ethanol exposure in mice than randomly constructed modules of the same size.

### Megamodules

Due to a large amount of redundancy between modules, our laboratory previously modified the EW-dmGWAS output by merging significant modules with greater than 80% overlap of genes[14]. Percent overlap represented the number of genes contained in both modules (for every possible pair) divided by the number of genes in the smaller module. The final merged modules are called “megamodules”. Megamodule scores were calculated using the EW-dmGWAS algorithm. Cytoscape was used to visualize the resulting megamodules, and an overrepresentation analysis was run to determine which megamodules had higher than expected number of genes significantly (p<1E-6) associated with alcohol consumption in the human GWAS dataset. These resulting overrepresentation p-values were used to select megamodules from each group for further functional enrichment analysis. Megamodules containing less than 10 genes were excluded from further analysis.

Gene ontology analysis of selected megamodules was then performed using ToppGene[16].

## Results

### Ethanol, GSK3β inhibition, and the interaction between the two differentially regulate genes

When compared to HV double controls, there were 345 significant differentially expressed genes (DEGs) in the HT group, 523 DEGs in the EV group, and 293 DEGs in the ET group within PFC. Within the NAc these treatment groups contained 556 (HT), 418 (EV), and 619 (ET) DEGs respectively. DEGs were determined using cutoffs of p<0.05 and LFC>0.1 or LFC<-0.1. Only a small percentage of genes are differentially expressed across multiple comparisons, suggesting separate and distinct populations of genes regulated by tideglusib treatment (HT), ethanol consumption (EV), and the interaction between ethanol and tideglusib (ET) (*Fig. 1*).

**Figure 1.**
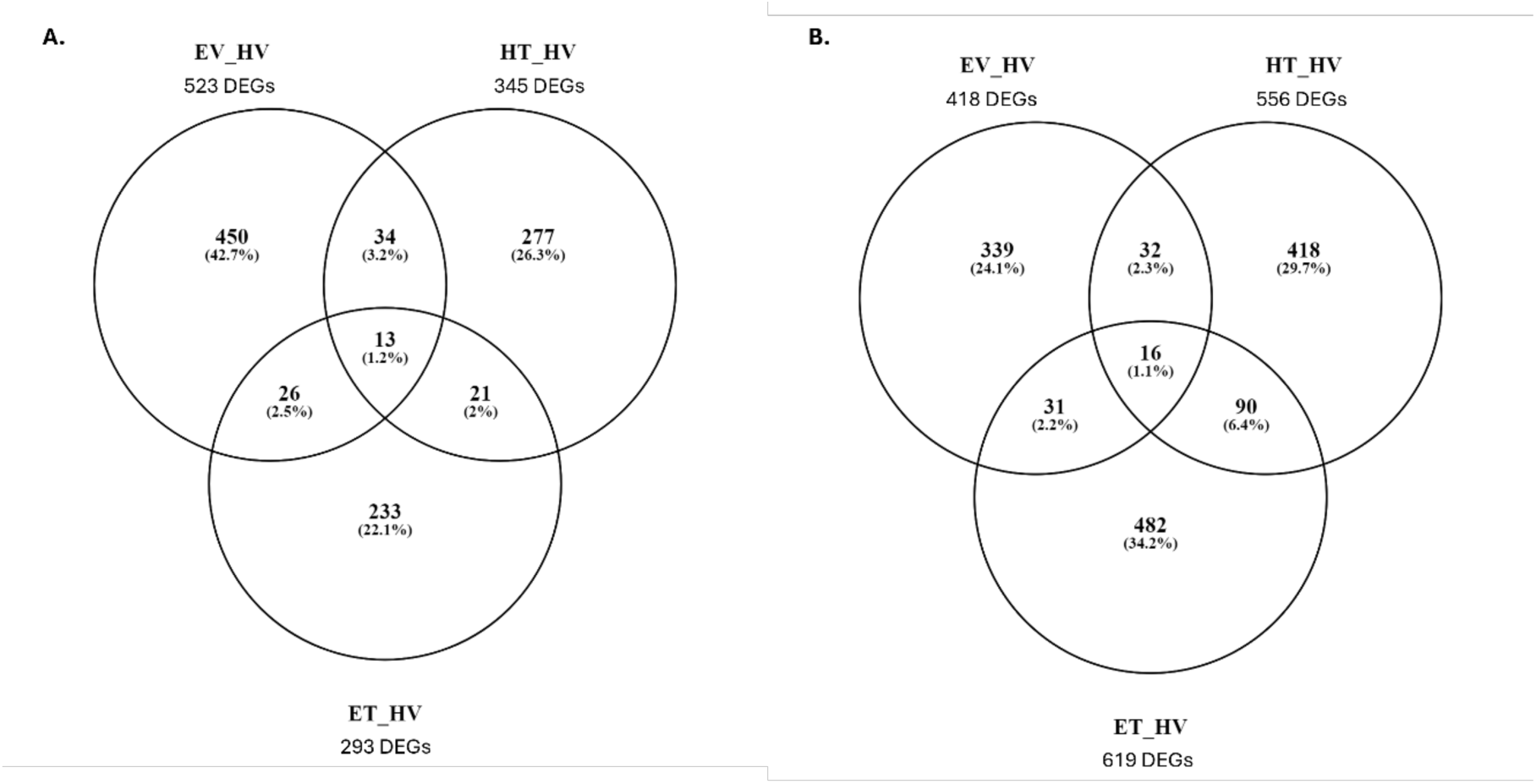
Ethanol, GSK3β inhibition, and the interaction of the two differentially regulate gene expression. **A.** PFC regulated genes. Ethanol regulated 523 DEGs (EV_HV), GSK3β inhibition regulates 345 DEGs (HT_EV), GSK3β inhibition in ethanol drinking mice regulates 293 DEGs (ET_HV) with low DEG overlap between groups. **B.** NAc regulated genes. Ethanol regulated 418 DEGs (EV_HV), GSK3β inhibition regulates 556 DEGs (HT_EV), GSK3β inhibition in ethanol drinking mice regulates 619 DEGs (ET_HV) with low DEG overlap between groups.

### GSK3β inhibition in ethanol-drinking mice mimics a water-drinking transcriptome within some gene clusters

Using the union of all DEGs identified across treatment groups within a given brain region, we used hierarchical cluster analysis to identify groups of co-regulated genes. Five clusters of DEGs were generated within each brain region using hierarchical clustering of z-scored transcript counts (*Fig. 2a*, *3a*). Both mPFC and NAC contained two clusters each where GSK3β inhibition appeared to partially reverse ethanol-induced differential gene expression, resulting in a transcript profile similar to both the HV and HT groups (PFC clusters 3 and 4: NAc clusters 1 and 5).

**Figure 2.**
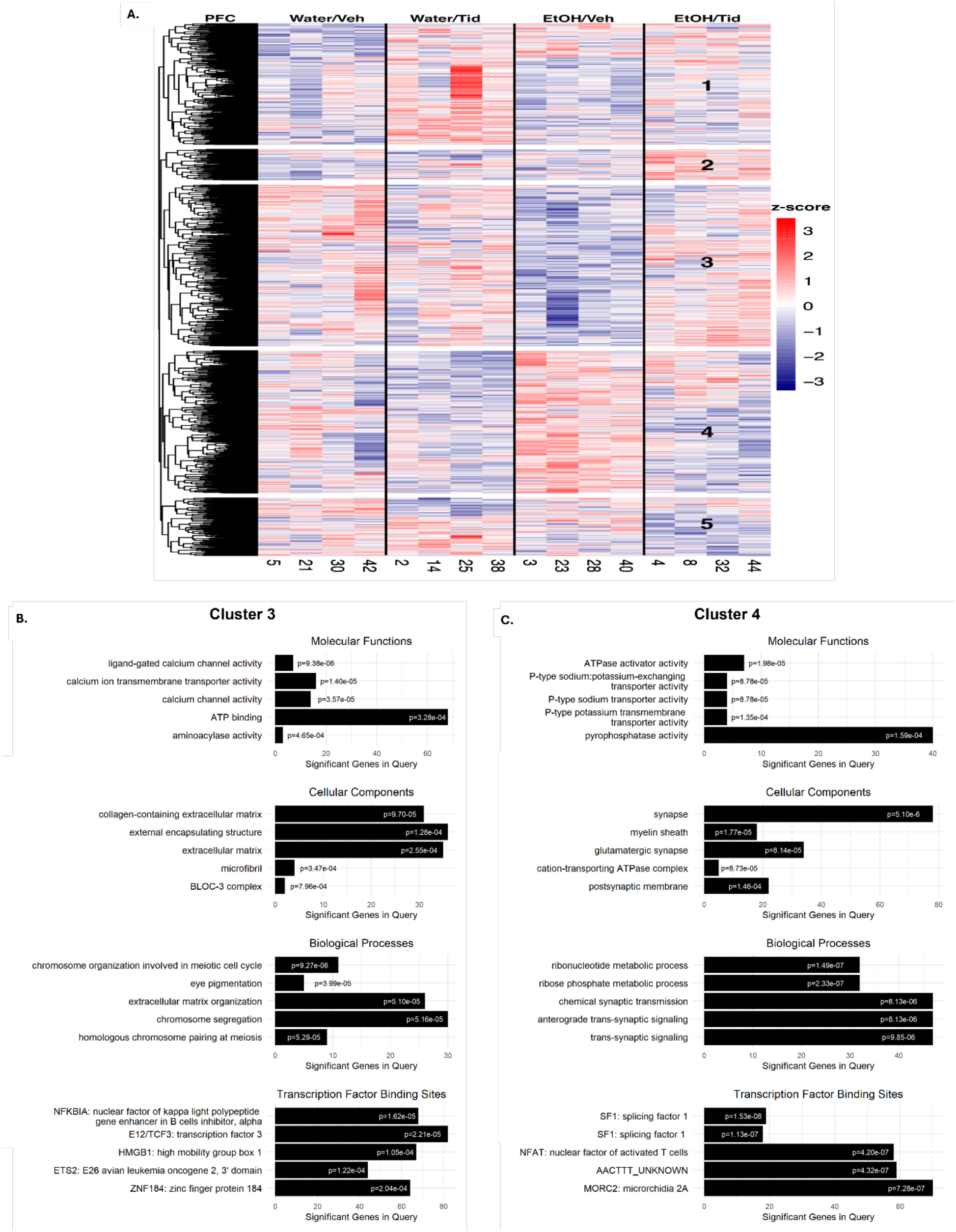
GSK3β inhibition reverses ethanol-induced transcriptional changes in specific gene clusters within mPFC. **A.** Hierarchical clustering map of genes differentially regulated across different treatment groups. **A-B.** GSK3β inhibition reverses ethanol-induced downregulation of genes in Cluster 3 and overexpression of genes in Cluster 4. Top five gene ontologies and their significant p-values are shown for molecular functions, cellular components biological processes, and transcription factor binding sites for each cluster.

### mPFC

In cluster 3 (*Fig. 2b*), gene ontology analysis identified 108 molecular functions significantly over-represented in categories which contained three or more genes within the genome that are known to play a role. The top three categories with the highest significance were all related to calcium channel activity, followed by ATP binding and aminoacylase activity to conclude the top five. Of the 65 cellular component categories identified, top results identified roles involved in extracellular matrix, external encapsulating structure, microfibrils, and lysosome-related organelles complex (BLOC-3). There were 464 biological processes category results. Top results again showed a role for genes in extracellular matrix, as well as chromosome organization. 310 transcription factor binding sites (TFBS) were identified, and top results were for NFKB inhibitor alpha (NFKBIA) and E12/TCF3, which is the TFBS for b-catenin in the canonical Wnt signaling pathway.

Cluster 4 (*Fig. 2c*) had 95 significantly overrepresented molecular functions, with three of the top five related to sodium/potassium transporter activity, and the others in the top five including ATPase activator and pyrophosphatase activity. There were 118 cellular component categories identified and the top five consisted of synapses, particularly glutamatergic synapses, postsynaptic membrane, myelin sheath, and ATPase complex. Of the 665 significant biological process categories, the top two were ribonucleotide and phosphate metabolic processes with the following three involved in synaptic transmission, particularly trans-synaptic signaling. 406 TFBS were identified as significant, and both top two results were for splicing factor 1 (SF1), followed by nuclear factor of activated T-cells (NFAT) and microrchidia 2A (MORC2).

### NAc

In cluster 1 (*Fig. 3b*), 71 significant molecular function categories were identified, and all the top five were related to regulating inhibitory signaling. The top two functions were for extracellularly glycine-gated channels, the next two most significant were for L-glutamate ligase activity, and the fifth most significant was for inhibitory extracellular ligand-gated monoatomic ion channel activity. Categories for 28 cellular components were identified, with top results being amylin receptor complex, microtubule organizing center, meiotic spindle, and microtubule cytoskeleton. Biological processes had 481 significant ontology results. Three of the top five results were for Wnt signaling, and the remaining two were hits for forebrain neurogenesis and positive regulation of signal transduction. TFBS results further implicated Wnt signaling in this cluster of genes. Of the 294 significant results, the top result was for E12/TCF3. Remaining categories in the top five results were for paired box 4 (PAX4), MYC-associated zinc finger protein (MAZ), hydroxysteroid 17-beta dehydrogenase 8 (HSD17B8), and an unknown AACTTT binding site.

**Figure 3.**
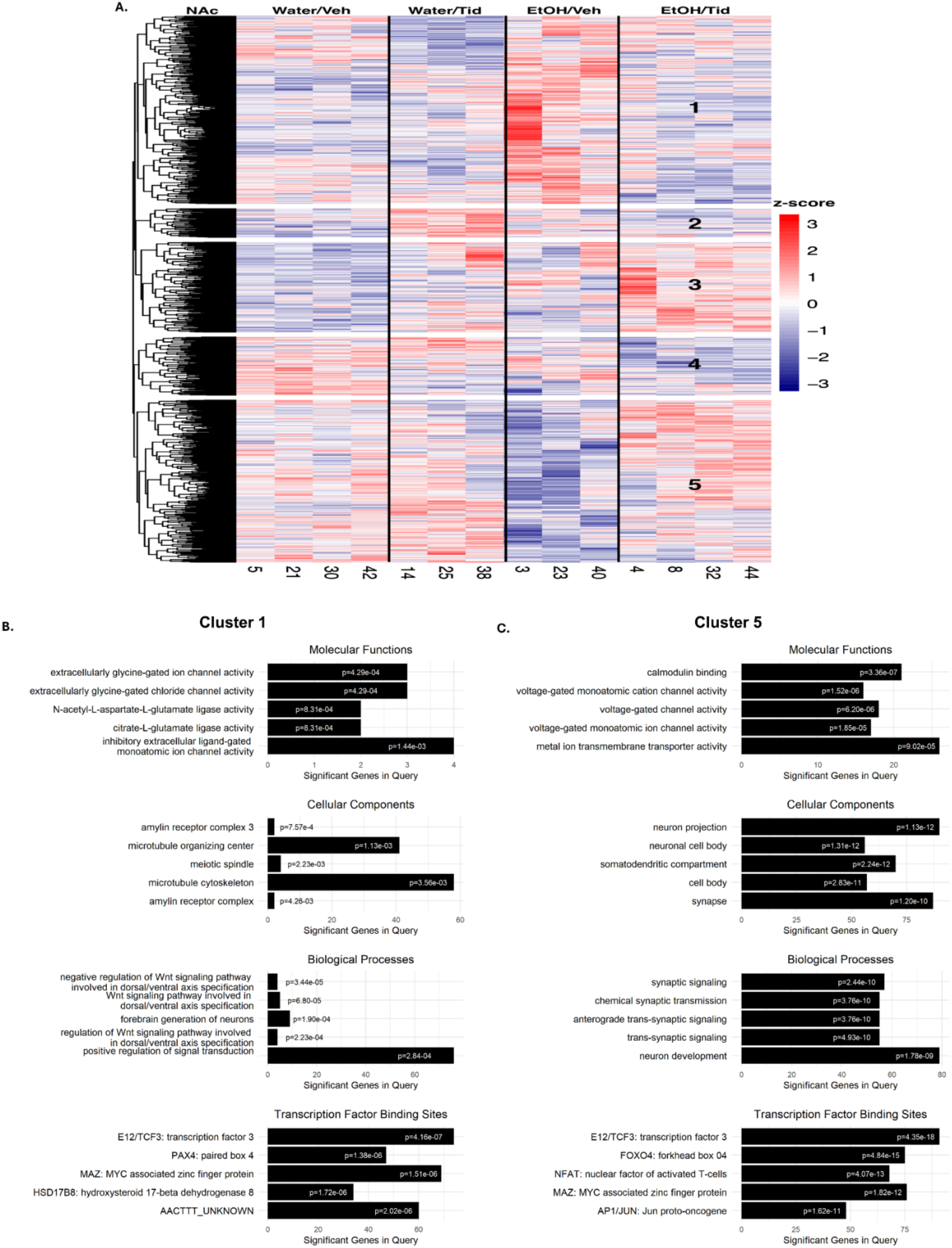
GSK3β inhibition reverses ethanol-induced transcriptional changes in specific gene clusters within NAc. **A.** Hierarchical clustering map of genes differentially regulated across different treatment groups. **A-B.** GSK3β inhibition reverses ethanol-induced increased expression of genes in Cluster 1 and decreased expression of genes in Cluster 5. Top five gene ontologies and their significant p-values are shown for molecular functions, cellular components biological processes, and transcription factor binding sites for each cluster.

Cluster 5 (*Fig. 3c*) had 126 molecular function ontology results, with three of the top results relating to voltage-gated channel activity and the others were calmodulin binding and metal ion transmembrane transporter activity. The top cellular component ontologies were neuron projection, neuron cell body, somatodendritic compartment, and synapse out of 129 significant results. Further implicating this cluster of genes in regulation of neurotransmission, out of 850 ontology results, the top four were for synaptic signaling and the fifth was for neuron development. Of the 376 TFBS identified, the top result was once again for E12/TCF3, followed by forkhead box 04 (FOXO4), NFAT, MAZ, and Jun proto-oncogene (AP1/JUN).

It is interesting that in the four clusters reported, E12/TCF3 appeared as either the first or second most significant TFBS within three. In PFC cluster 4, where we do not see E12/TCF3, the top TFBS results are both splicing factor 1 (SF1) which has also been suggested to be regulated by β-catenin[22].

### EW-dmGWAS identifies gene networks regulated by chronic ethanol and the interaction of ethanol*GSK3β inhibition

DEG analysis allowed for identification of genes regulated by ethanol, GSK3β inhibition via tideglusib, and through the interaction of the two. Using human EW-dmGWAS, we can combine these results with human GWAS data to identify expressional changes from our manipulations specifically within genes known to be associated with problematic alcohol consumption in humans. By further incorporating known PPI, expressional networks of DEGs with known correlations to AUD can be identified. We found three networks regulated by chronic ethanol consumption (EV_HV) and two networks regulated by ethanol plus tideglusib (ET_HV) in the mPFC compared to water drinking animals and identified the hub gene of each network. Interestingly, tideglusib treatment alone did not produce any significant results. As these analyses utilize an overrepresentation analysis of DEGs specifically within known AUD-correlated GWAS results, it can be concluded GSK3β inhibition in the absence of ethanol does not elicit expressional changes within AUD- associated genes. A summary of significant PFC EW-dmGWAS results is in *Table 1*.

**Table 1.** Significant expression networks identified by EW-dmGWAS ***Table key:*** *Comparison, treatment groups being compared; Megamod, name of expression network;Size (genes), number of genes in network determined by expression values from RNAseq and known protein-protein interactions; GWAS hits in MegaMod, number of genes from RNAseq significant in alcohol consumption GWAS; p-value, significance value of network based on overrepresentation of network genes significant within GWAS; Overrep genes, genes with significant GWAS hits; Odds Ratio, strength of correlation between genes from RNAseq and overrepresentation in GWAS; Hub gene, central gene of the expression network; Edges, number of protein-protein interactions between hub gene and other genes in network based on correlated expressional changes between treatment groups and known protein-protein interactions*.

| Comparison | MegaMod | Size (genes) | GWAS hits in MegaMod | p-value | Overrep genes | Odds Ratio | Hub gene | Edges |
| --- | --- | --- | --- | --- | --- | --- | --- | --- |
| EV vs HV | navyblue | 17 | 6 | 6.75E-08 | CBX5<br>EP300<br>MAPT<br>PPM1G<br>PSMA3<br>PSMC3 | 40.233 | PSMA1 | 10 |
| EV vs HV | goldenrod_Two | 13 | 5 | 5.42E-07 | ARPC1B<br>ATXN2L<br>MAPT<br>RANGAP1<br>RUNX1T1 | 45.926 | PRKACB | 5 |
| EV vs HV | purple | 13 | 5 | 5.42E-07 | ARPC1A<br>CAD<br>CLN3<br>TRMT10A<br>TUFM | 45.926 | XPO1 | 4 |
| ET vs HV | lawngreen | 20 | 5 | 6.04E-06 | CBX5<br>EP300<br>GSE1<br>MAPT<br>PSMC3 | 24.485 | VHL | 11 |
| ET vs HV | seashell | 16 | 5 | 1.78E-06 | FAF1<br>GTF3C2<br>MAPT<br>PPM1G<br>SPI1 | 33.396 | N/A | N/A |

### Chronic ethanol regulated expression networks

Three networks were identified as regulated by chronic ethanol in mPFC. In order of significance, regarding over- representation of GWAS results, they are navyblue (p=6.75E-08), goldenrod_two (p= 5.42E- 07), and purple (p=5.42E-07).

There are 17 genes in the navyblue network, with six identified as associated with problematic alcohol consumption in GWAS results. The hub gene within this network is proteosome subunit alpha 1 (PSMA1), a deubiquitinating protein that has been shown to regulate Akt and β-catenin levels and promotes cellular proliferation[23] (*Fig. 4*). Notably, a multitude of other proteosome subunits appear in this network and strongly implicate chronic ethanol consumption in proteosome regulation.

**Figure 4.**
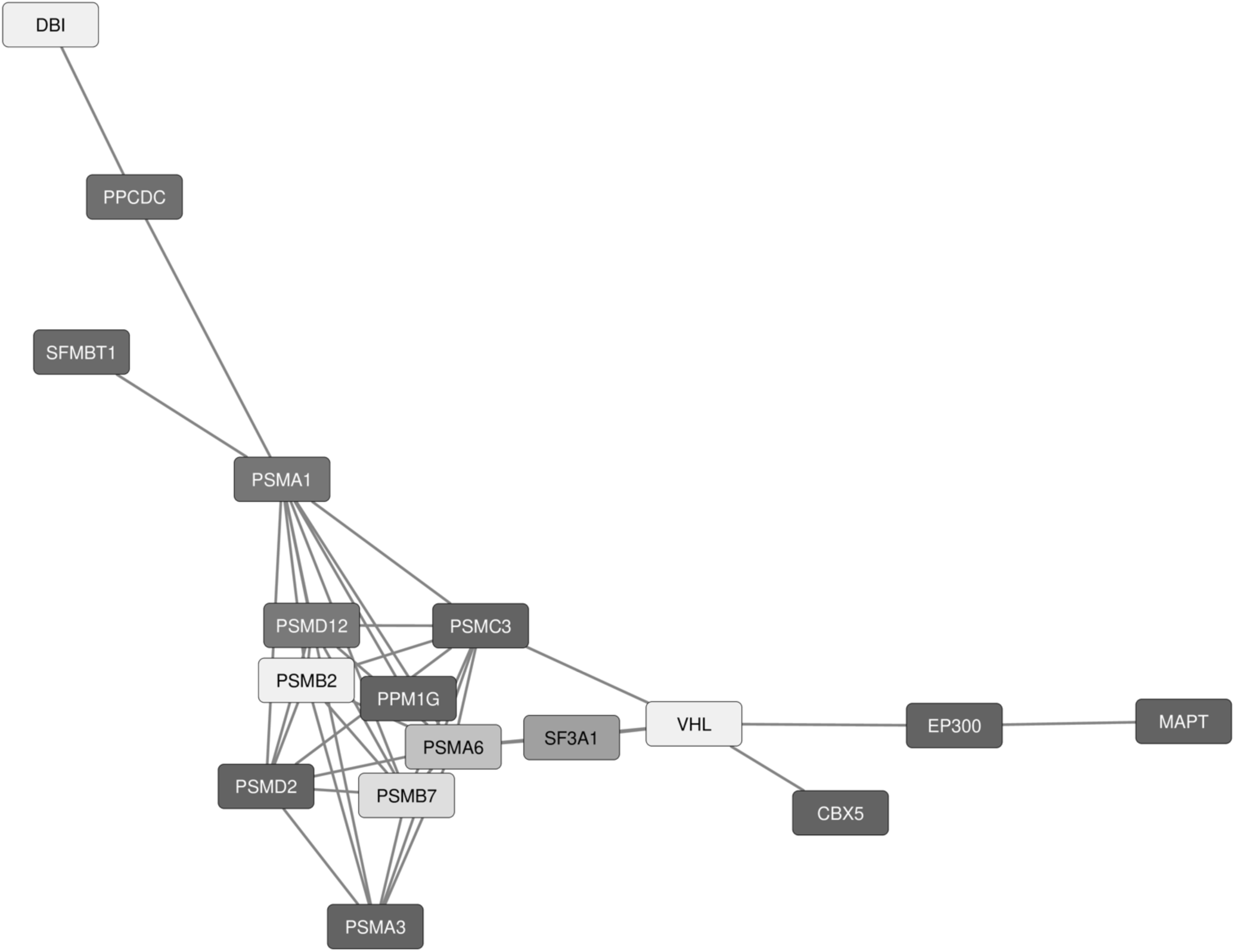
Navyblue gene expression network regulated by chronic ethanol consumption (EV_HV) (6.75E-08). Network contains 17 genes from PFC RNAseq analysis, six of which are significant in GWAS (CBX5, EP300, MAPT, PPM1G, PSMA3, PSMC3). Central hub gene is PSMA1 which has 10 edges connecting to other genes. Darker colors correlate to greater GWAS over-representation significance; shorter edges correlate to greater edge weights and more significant PPI.

Gene ontology analysis identified 44 significant molecular function results, the first of which was NFKB binding followed by transcription factor binding, benzodiazepine binding, and proteosome-activating activity (*Fig. 5a*). The cellular and molecular results also implicated this pathway in proteosome regulation. Of the 44 cellular component results, top hits were for proteosome complex, peptidase complex, and catalytic complex (*Fig. 5b*). Biological processes had 299 significant ontology results, and all the top five were involved in proteosome and ubiquitin-dependent catabolic processes (*Fig. 5c*). TFBS returned 72 significant ontology results, and all the top three were related to nuclear factor, erythroid derived 2 (NFE2), a known immune regulator[24], followed by ELK1 member of the ETS oncogene family, and YY1 transcription factor (*Fig. 5d*). Given PSMA1 is central to this network, functional proteosome involvement is unsurprising. Likewise, effects on transcription factor binding and immune reactivity logically track given the actions of PSMA1 in cancer cell proliferation and its ability to regulate β-catenin[23, 25, 26]. Effects on benzodiazepine receptor binding is interesting given the similar effects at the GABA_A_ receptor between benzodiazepines and ethanol and the development of cross-tolerance between the two drugs[27, 28]. However, it is important to note this ontology result has a single DEG contributing to this category. Low query hits are not unexpected given the relatively few numbers of genes within each network, however it does highlight the need for caution when interpreting the ontology results. Results with a higher number of query hits may thus constitute more accurate functions of our networks.

**Figure 5.**
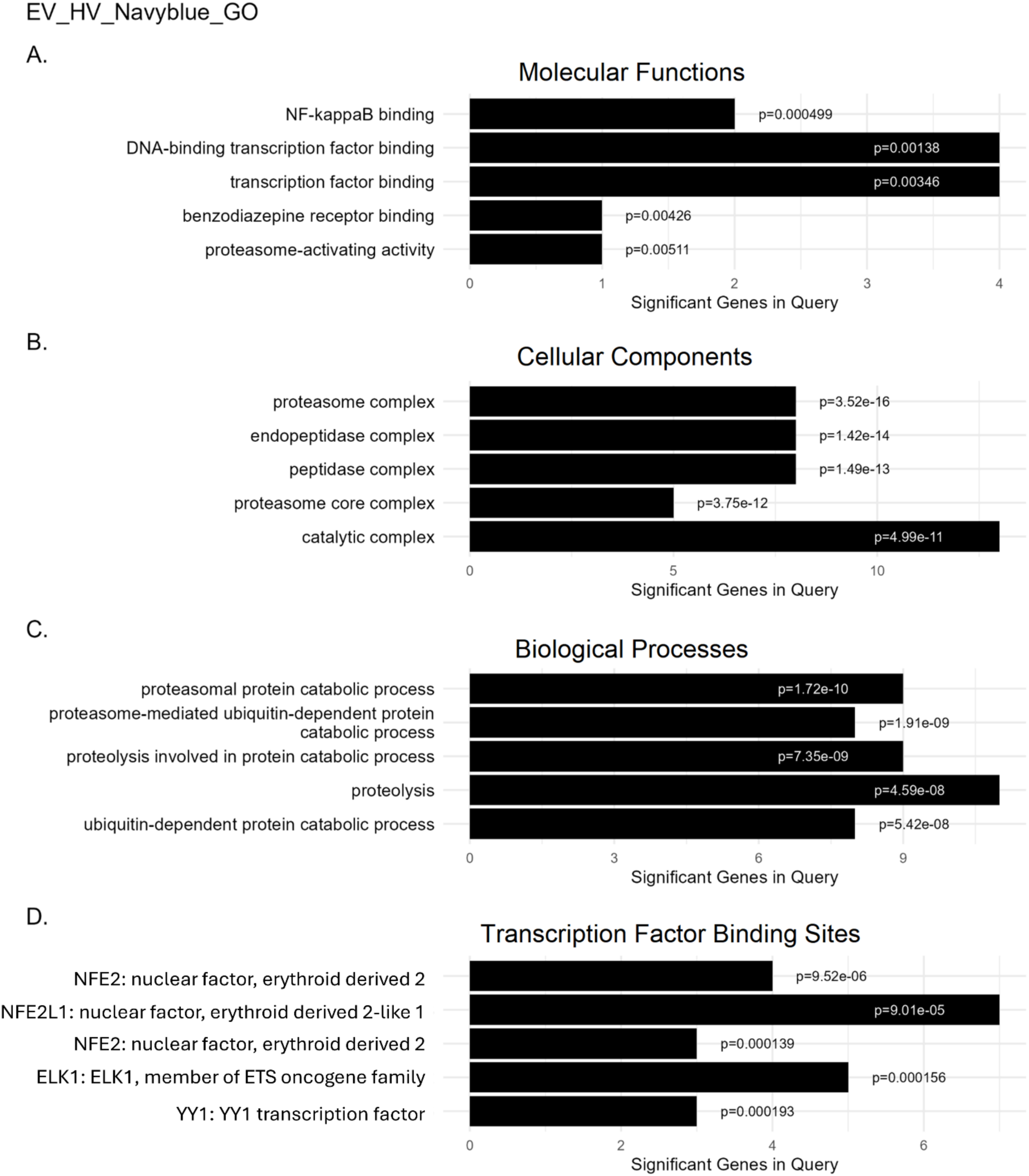
The top five gene ontologies for the navyblue expression network and significant p-values are shown. **A** Molecular functions, **B** Cellular components, **C** Biological processes, and **D** Transcription factor binding sites. Gene network is functionally overrepresented with genes involved in proteosome and immune reactivity.

The goldenrod_two network is comprised of 13 genes, and 5 appeared in GWAS results. The hub gene was protein kinase cAMP-activated catalytic subunit beta (PRKACB). PRKACB is the catalytic subunit of protein kinase A (PKA), a serine/threonine kinase which mediates signaling through cyclic AMP (cAMP)[29]. Alcohol induces subcellular translocation of the PKA catalytic subunit into the nucleus[30], and decreases in PKA levels or PKA inhibition has been shown to decrease alcohol consumption in rodents[31, 32]. PKA is also known to phosphorylate GSK3β[33]. Here we show an ethanol-regulated PKA-centric network (*Fig. 6*).

**Figure 6.**
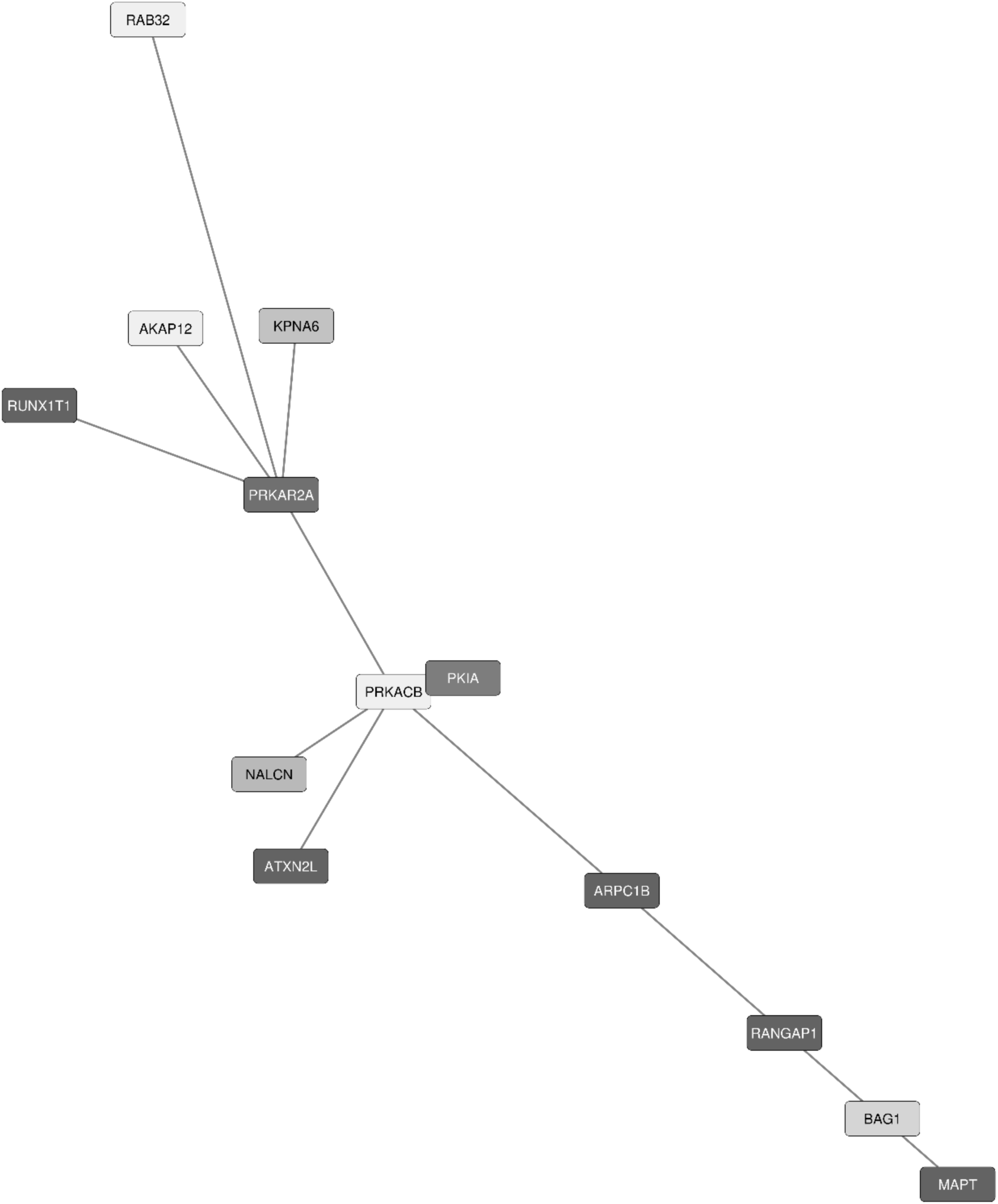
Goldenrod_two gene expression network regulated by chronic ethanol consumption (EV_HV) (5.42E-07). Network contains 13 genes from RNAseq analysis, five of which are significant in GWAS (ARPC1B, ATXN2L, MAPT, RANGAP1, RUNX1T1). Central hub gene is PRKACB which has 5 edges connecting to other genes. Darker colors correlate to greater GWAS overrepresentation significance; shorter edges correlate to greater edge weights and more significant PPI.

Gene ontology analysis identified 53 molecular functions, 53 cellular components, and 226 biological processes. Nearly every result in the top five of each category dealt with PKA binding or activity with some additional results for ubiquitin protein ligase binding, cytoplasm, and negative regulation of phosphorylation (*Fig. 7a-c*). TFBS had 36significant results. The top result was for an unknown binding site, but the second most significant hit was for TEA domain family member 2 (ETF), a known mediator of proliferation-dependent differential gene expression through cytokine-independent MAPK signaling[34]. Other top TFBS results were for multiple zinc finger proteins (ZNFs) and nuclear respiratory factor 1 (NRF1 (*Fig. 7d*). NRF1 deletion has been linked to impaired proteosome function and neurodegeneration[35], and overexpression has been shown to prevent dopaminergic neuron degeneration in a mouse model of Parkinson’s disease[36]. These effects on the proteosome are thought to be through actions on deubiquitinating proteins[37] as was seen in the previous expression network.

**Figure 7.**
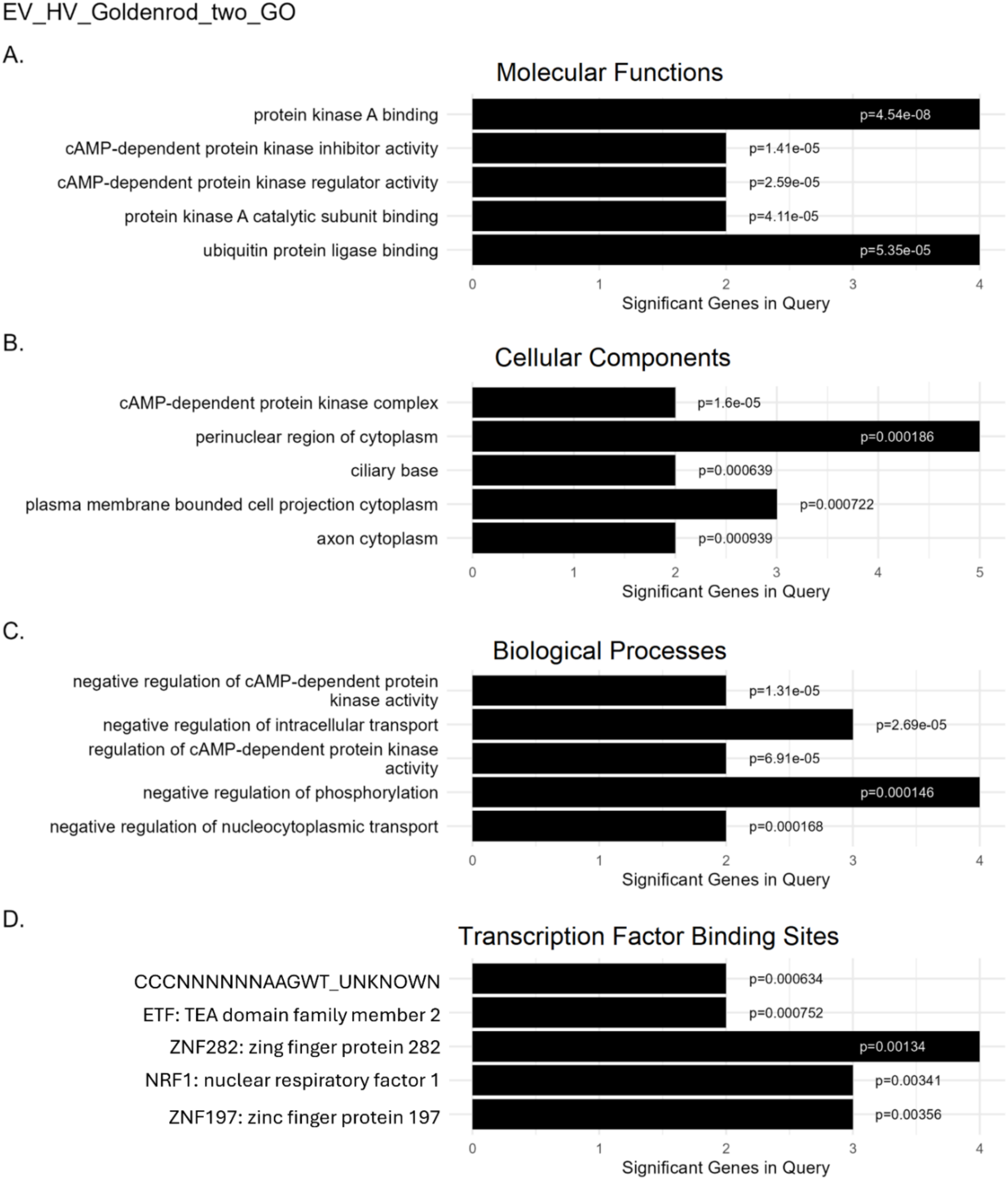
The top five gene ontologies for the goldenrod_two expression network and significant p-values are shown. **A** Molecular functions, **B** Cellular components, **C** Biological processes, and **D** Transcription factor binding sites. Gene network is functionally overrepresented with genes involved in PKA binding/activity.

The purple expression network is comprised of 13 genes, five of which are significant in GWAS results. The central member of this network is exportin 1 (XPO1) (*Fig. 8*). Originally named chromosomal region maintenance 1 (CRM1), XPO1 regulates dynamic response to genotoxic stress and enables nuclear export from the nucleus[38]. Additionally, Akt3 has been shown to regulate XPO1-dependent nuclear export in an Akt1-independent manner[39].

**Figure 8.**
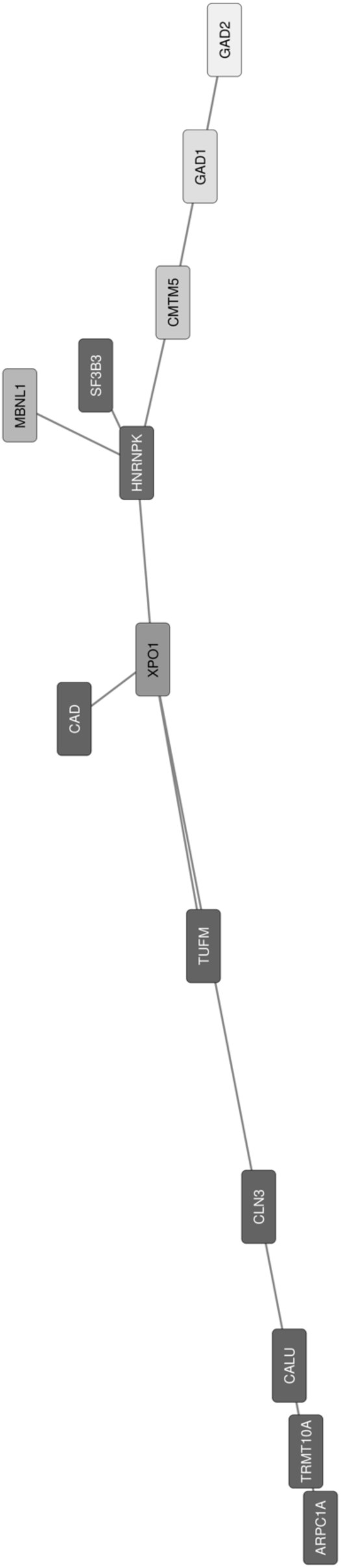
Purple gene expression network regulated by chronic ethanol consumption (EV_HV) (5.42E-07). Network contains 13 genes from RNAseq analysis, five of which are significant in GWAS (ARPC1B, CAD, CLN3, TRMT10A, TUFM). Central hub gene is XPO1 which has 4 edges connecting to other genes. Darker colors correlate to greater GWAS over- representation significance; shorter edges correlate to greater edge weights and more significant PPI.

This gene network had 37 significant ontology results for molecular functions. Top results were for amino acid binding, glutamate binding, carboxy-lyase activity, pyridoxal phosphate binding, and vitamin B6 binding (*Fig. 9a*). Of the 47 cellular components identified, the top three all involved clathrin-sculpted vesicles, followed by presynapse and inhibitory synapse (*Fig. 9b*). These results implicate this network in neurotransmission, specifically at the step of presynaptic endocytosis. Clathrin-mediated endocytosis (CME) is the main mechanism for internalization into the cell and plays a pivotal role in synaptic vesicle recycling[40]. Clathrin polymerizes to form a clathrin-coated vesicle which is then separated from the plasma membrane via scission by dynamin[41]. Continued neurotransmission following an initial transmitter release is dependent on CME. Interestingly, GSK3β is a known facilitator of activity-dependent bulk endocytosis (ADBE), where it rephosphorylates dynamin I to prepare it for the next round of ADBE[42]. Further supporting a role for this network in presynaptic neurotransmission are the biological process results. There were 225 ontology results for this category, and all the top five were involved with amino acid biosynthesis, specifically with glutamine and glutamate metabolism (*Fig. 9c*). Given the cycle of glutamine-glutamate/GABA, it seems likely this pathway may be responsible for shifts in excitatory/inhibitory balance following chronic ethanol. In this cycle, glutamine is converted to glutamate within excitatory glutamatergic neurons, and glutamate is further metabolized to GABA within inhibitory GABAergic cells.

**Figure 9.**
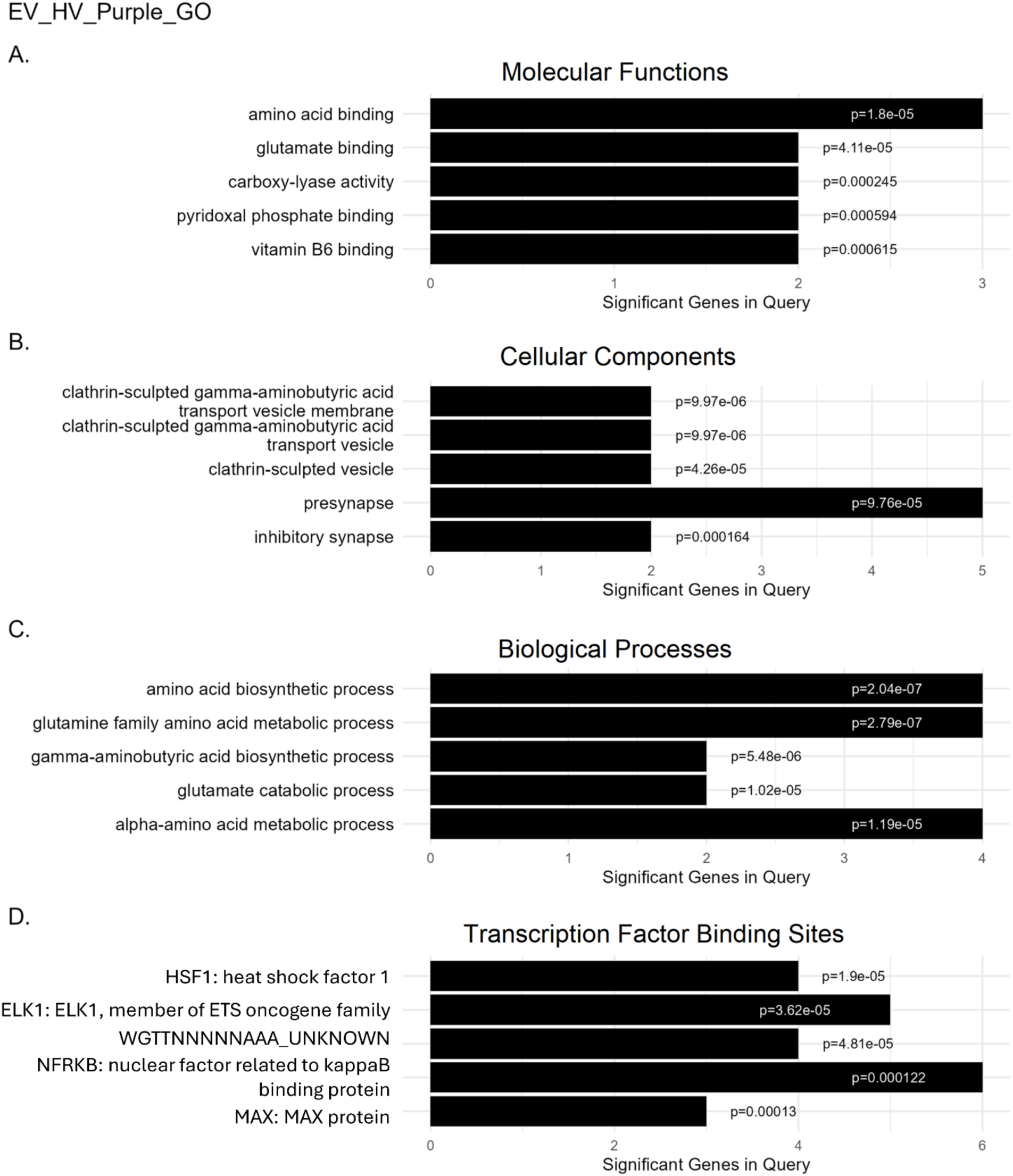
The top five gene ontologies for the purple expression network and significant p-values are shown. **A** Molecular functions, **B** Cellular components, **C** Biological processes, and **D** Transcription factor binding sites. Gene network is functionally overrepresented with genes involved in presynaptic signaling.

Released glutamate/GABA is then accumulated in astrocytes where it is converted back to glutamine and released to reaccumulate in neurons, continuing the cycle[43]. In combination, shifts within this cycle plus effects on vesicle recycling would result in altered excitatory/inhibitory synaptic transmission. Additionally, 99 TFBS were identified in our functional analysis. The top TFBS was heat shock factor 1 (HSF1), followed by ELK1, nuclear factor related to kappaB binding protein (NFRKB), and MYC associated factor X (MAX) (*Fig. 9d*). HSF1 is a stress-responsive transcription factor which promotes cell survival. Ethanol induces HSF1 nuclear translocation, and HSF1 activity regulates NF-kappaB[44, 45]. Additionally, HSF1 has been shown to bind to *Bdnf* promoters and induce BDNF expression[46]. Interestingly, BDNF is documented to interfere with GSK3β rephosphorylation of dynamin I, inhibiting ADBE[47] further implicating this network in ethanol-induced effects on neurotransmission.

### Chronic ethanol plus tideglusib regulated expression networks

Two networks were identified as regulated by chronic ethanol plus tideglsuib in mPFC. In order of significance, they are lawngreen (p=6.04E-06) and seashell (p=1.78E-06).

The lawngreen network contains 20 genes, five of which were significant in GWAS, and has von Hippel-Lindau tumor suppressor (VHL) as the central member (*Fig. 10*). VHL is a tumor suppressor gene which encodes two gene products: pVHL_30_ and pVHL_19_ [48]. pVHL_30_ functionally associates with microtubules and has been identified as a target of GSK3[48]. VHL has also been shown to work in tandem with GSK3β to maintain cellular cilia through these microtubule interactions[49].

**Figure 10.**
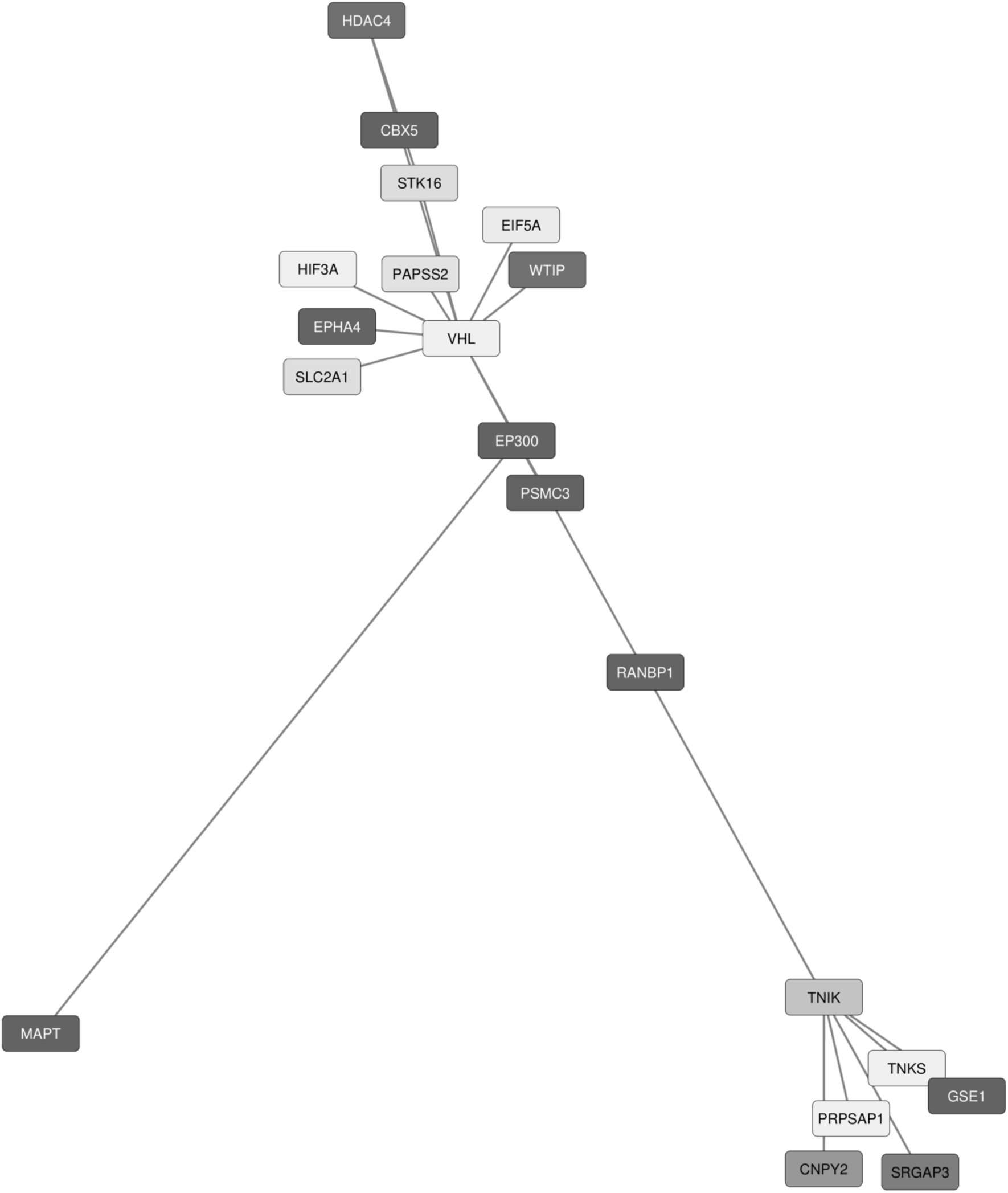
Lawngreen gene expression network regulated by GSK3β inhibition in animals with chronic ethanol consumption (ET_HV) (6.04E-06). Network contains 20 genes from RNAseq analysis, five of which are significant in GWAS (CBX5, EP300, GSE1, MAPT, PSMC3). Central hub gene is VHL which has 11 edges connecting to other genes. Darker colors correlate to greater GWAS over-representation significance; shorter edges correlate to greater edge weights and more significant PPI.

Of the 88 detected molecular functions, top results were for kinase binding and transferase activity, with the remaining of the top five all relating to transcriptional regulation (*Fig. 11a*). Cellular components returned 69 significant ontology results. The top result again related to transcription regulation, followed by nuclear pore, axonal growth cone, postsynaptic density, and asymmetric synapse (*Fig. 11b*). Given the roles of GSK3β in both microtubule and postsynaptic density 95 (PSD-95) stabilization, these results are consistent with GSK3β inhibition. GSK3β has known functions on microtubules outside of the previously mentioned interaction with VHL, where in disease free conditions, GSK3β phosphorylates tau, which in turn promotes tubulin assembly into microtubules. In tau pathology, GSK3β hyperphosphorylates tau, and this GSK3β-induced hyperphosphorylation is currently under investigation for treatment of Alzheimer’s disorder[50, 51]. Additionally, GSK3β phosphorylates PSD-95, destabilizing the anchoring protein, and resulting in internalization of glutamatergic receptors AMPA and NMDA[52, 53]. 594 biological process results were also identified, and the top four results were all related to hypoxia and response to oxygen levels. The fifth result was positive regulation of cell projection organization, more evidence of this pathway’s involvement in microtubules (*Fig. 11c*). The central member of this network, VHL, promotes degradation of hypoxia-inducible factor 1 (HIF-1). Interestingly, GSK3β overexpression has also been shown to reduce HIF-1a in a VHL-independent manner through phosphorylation-induced targeting of HIF-1a to the proteosome[54].

**Figure 11.**
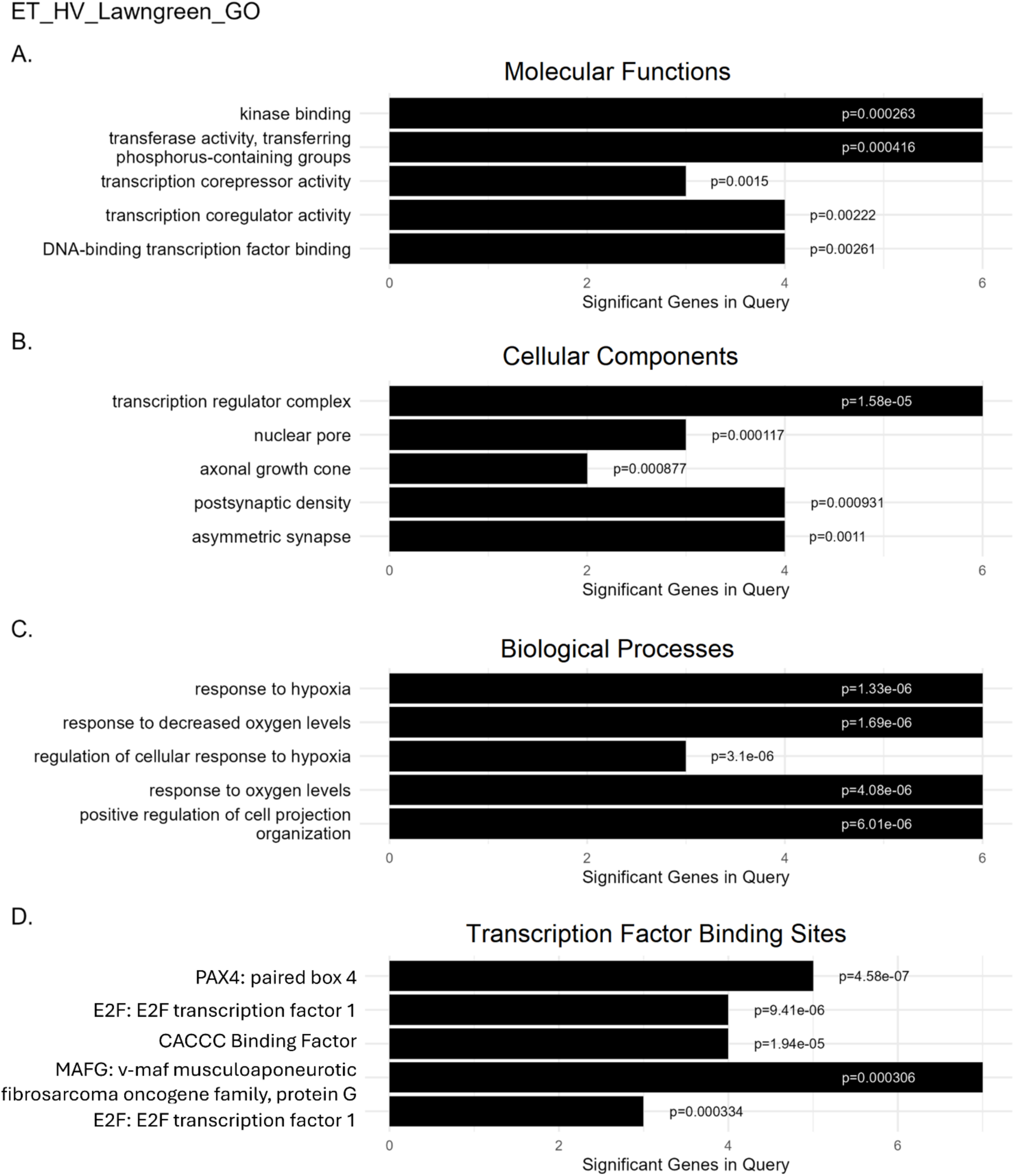
The top five gene ontologies for the lawngreen expression network and significant p-values are shown. **A** Molecular functions, **B** Cellular components, **C** Biological processes, and **D** Transcription factor binding sites. Gene network is functionally overrepresented with genes involved in proteosome and immune reactivity

Further supporting GSK3β mediation of response to oxygen levels is the top TFBS result, paired box 4 (PAX4). Of the 133 identified TFBS, PAX4 was the most significant and has been shown to activate genes encoding for proteosome subunits[55]. Additionally, VHL deletion has been shown to decrease PAX4 expression [56]. Additional top TFBS were E2F transcription factor (E2F) and v-maf musculoaponeurotic fibrosarcoma oncogene family, protein G (MAFG) (*Fig. 11d*).

Macrophage-activating factor (Maf) is a family of oncogenes, and MAFG has been shown to dimerize with the large subunit of NFE2[57]. The NFE2 TFBS was differentially regulated in the navyblue network regulated by chronic ethanol consumption, and the lawngreen network shares four of its five GWAS significant genes with navyblue (CBX5, EP300, MAPT, PSMC3). Additionally, of those four, only microtubule associated protein tau (MAPT) is expressed in different directions between the groups compared to control HV animals (lower expression in EV mice, higher expression in ET mice). The overlap of genes between these two networks and the similarity in expressional changes within these genes between treatment groups suggests the lawngreen network is regulated by ethanol rather than the ethanol*GSK3β inhibition interaction. It thus seems, at least in this expression network, GSK3β inhibition is unable to overcome the ethanol-induced changes in gene expression.

Conversely, the seashell network contains functional categories unique to the ethanol*GSK3β interaction group. This network consists of 16 genes, five of which are significant in GWAS results. This network is unique amongst our results in that there is no central hub gene, instead consisting of multiple genes with large edge-weights, creating a significant, but far spread network (*Fig. 12*). Gene ontology analysis returned significant results for 57 molecular functions, 26 cellular components, and 352 biological processes. The top five of each category was overrepresented with functions related to transcriptional regulation (*Fig. 13a-c*). 55 TFBS were significant in this network, with the top results being zinc finger E-box binding homeobox 1 (ZEB1), signal transducer and activator of transcription 5A (STAT5A), E12/TCF3, and LIM homeobox protein 2 (LHX2) (*Fig. 13d*). As previously mentioned, E12/TCF3 is a TFBS for β-catenin within the canonical Wnt signaling pathway[58], however each of the other TFBS identified has some involvement in Wnt signaling. ZEB1 represses Wnt inhibitory protein 1 (WIF1)[59], and in turn WIF1 signaling decreases ZEB1 expression[60]. ZEB1 has also been shown to activate Wnt signaling by upregulating β-catenin[61]. LHX2 has also been implicated in canonical Wnt-β-catenin signaling. Deleting *Lhx2* in cortical progenitor cells prevents Wnt-β-catenin signaling from maintaining cortical progenitor cell proliferation, resulting in fewer neurons across all cortical layers and smaller cortex volume[62]. Additionally, STAT5 has been linked to Wnt signaling through non-canonical, β-catenin-independent signaling. GSK3β phosphorylates STAT5, activating it and allowing STAT5 to bind to the promoter of the Wnt antagonist, secreted frizzled-related proteins (*Sfrp*)[63]. Together, there is clear evidence tideglusib treatment in ethanol drinking mice is modulating an expression network highly connected to Wnt signaling pathways, further supporting the analysis above.

**Figure 12.**
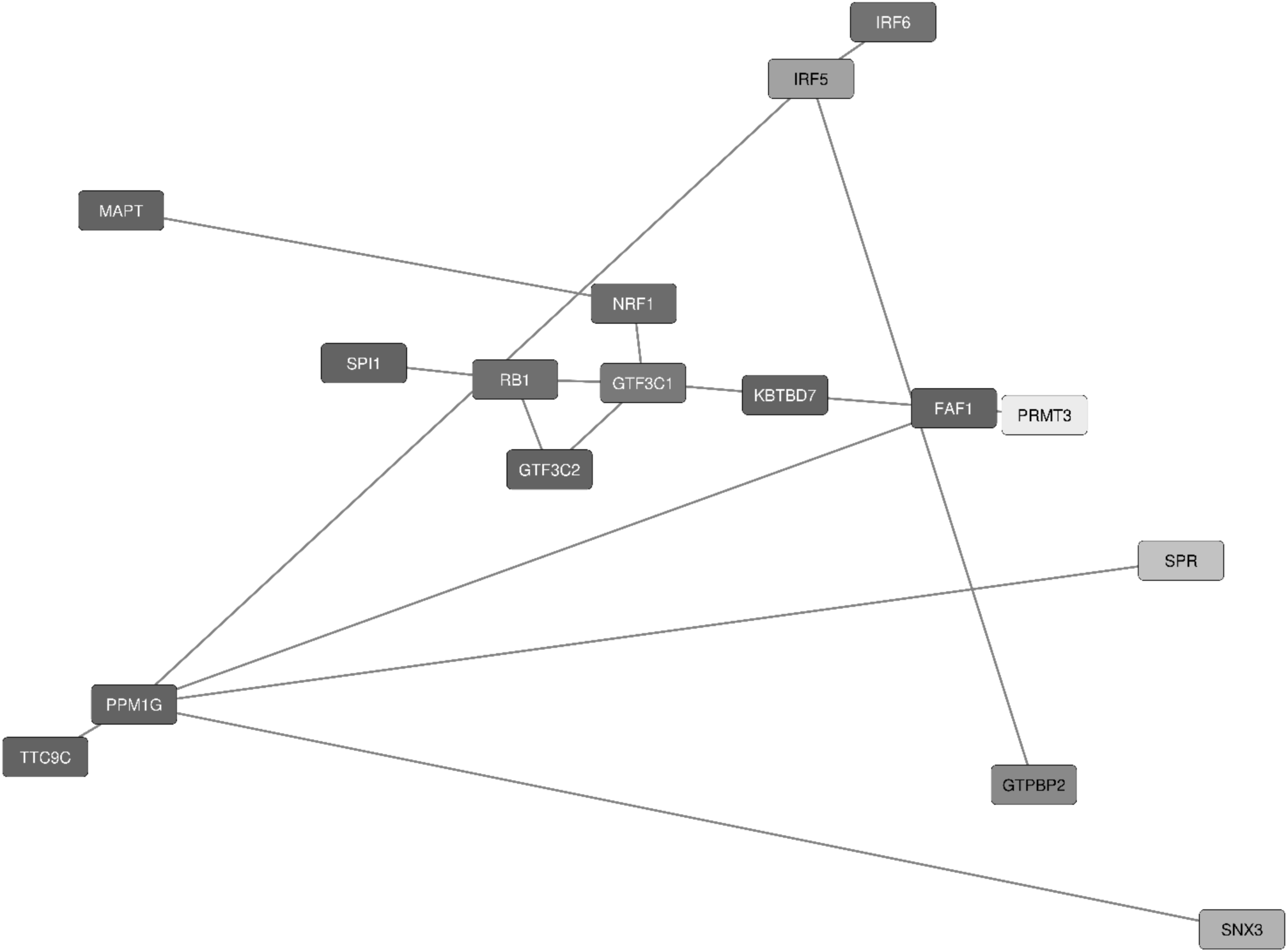
Seashell gene expression network regulated by GSK3β inhibition in animals with chronic ethanol consumption (ET_HV) (1.78E-06). Network contains 16 genes from RNAseq analysis, five of which are significant in GWAS (FAF1, GTF3C2, MAPT, PPM1G, SPI1). There is no central hub. Darker colors correlate to greater GWAS over-representation significance; shorter edges correlate to greater edge weights and more significant PPI.

**Figure 13.**
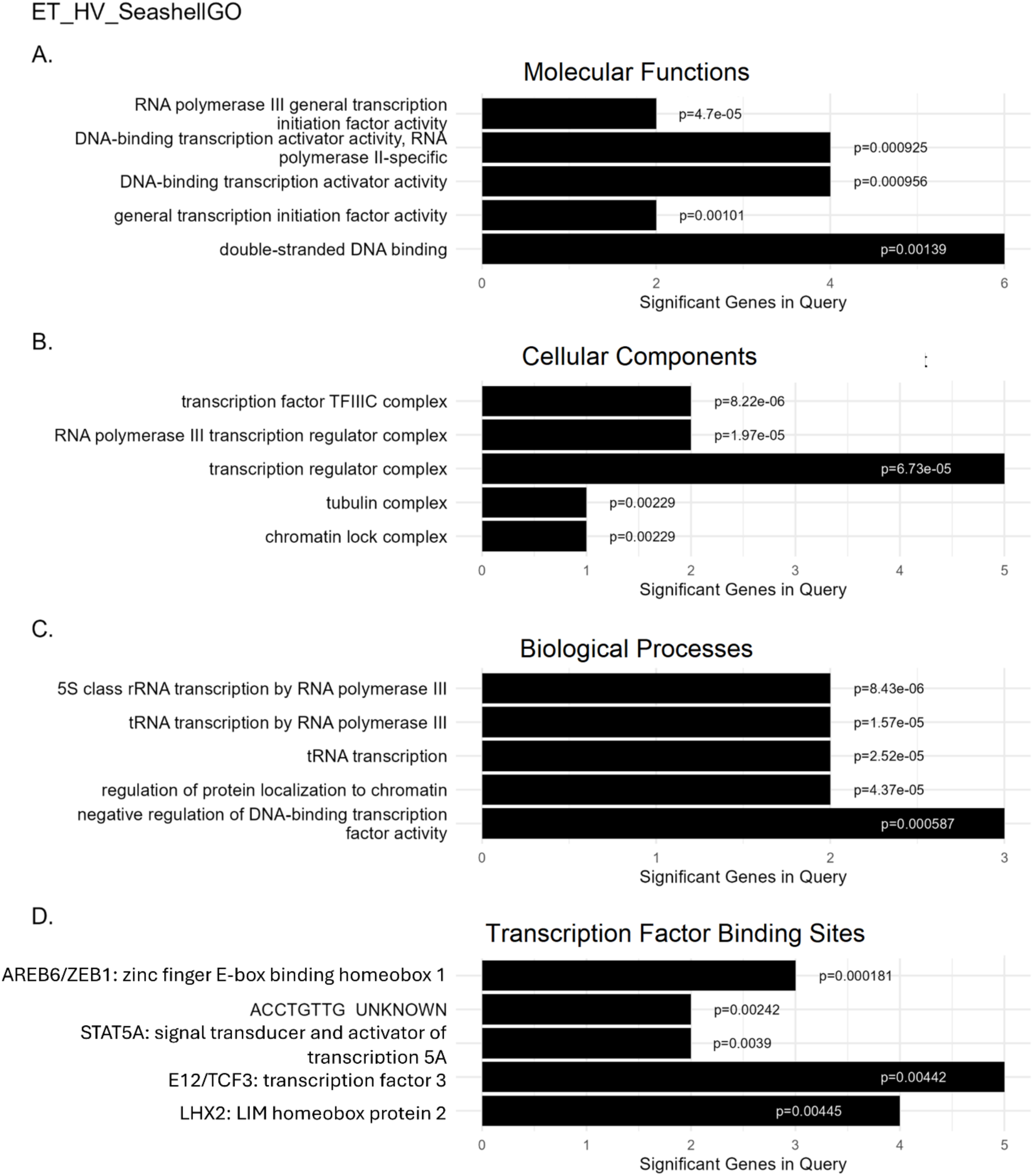
The top five gene ontologies for the seashell expression network and significant p-values are shown. **A** Molecular functions, **B** Cellular components, **C** Biological processes, and **D** Transcription factor binding sites. Gene network is functionally overrepresented with genes involved in transcription.

## Discussion

Here we have shown GSK3β inhibition partially reverses ethanol-induced gene expression changes to replicate an ethanol-naïve transcriptome within specific clusters of genes in both the mPFC and NAc. These clusters were overrepresented for genes implicated in the extracellular matrix, voltage-gated calcium channels, extracellular ligand gated channels (specifically glycine), glutamate metabolism, neuronal projections, and synapses/synaptic signaling, suggesting GSK3β inhibition is specifically acting to ameliorate or prevent the maladaptive effects on neurotransmission accompanying chronic ethanol consumption. GSK3β activity has previously been shown to regulate synaptic signaling both pre- and post-synaptically. GSK3β inhibits presynaptic neurotransmitter release through phosphorylation of voltage-gated calcium channels[64] and promotes ADBE through phosphorylation of dynamin 1[42]. Additionally, GSK3β destabilizes both excitatory and inhibitory post synaptic receptors by phosphorylating PSD-95 and gephyrin, resulting in internalization of AMPA/NMDA and GABA receptors[52, 53, 65, 66].

Additionally, our transcriptomic cluster analysis revealed a shift of gene expression in Wnt signaling related genes following tideglusib administration in ethanol drinking animals towards a water drinking expression profile. This was true in both brain regions, though there seems to be a stronger effect in NAc where top biological processes ontology was overwhelmingly Wnt related. Nonetheless, ontology results for the canonical TFBS for β- catenin, E12/TCF3, were significantly different in both PFC and NAc, as were other TFBS with known Wnt signaling interactions.

While actions of GSK3β on both neurotransmission and Wnt signaling have been firmly established, this is the first study to show GSK3β inhibition can specifically potentially reverse chronic ethanol-induced actions on these functions. Additionally, expression network analysis further supported a reversal of maladaptive synaptic signaling following GSK3β inhibition. We identified three EW-dmGWAS networks which contained known gene associations to problematic alcohol use that were regulated by chronic ethanol consumption in our animals. The dominant functions of these networks were proteosome activity/immune reactivity (navyblue), PKA binding/activity (goldenrod_two), and presynaptic signaling (purple). Interestingly, tideglusib treatment failed to counteract expressional changes within the proteosome/immune network, revealing a similar expression network in the ethanol*GSK3β inhibition animals (lawngreen) which contained an overlap of genes with navyblue which were similarly regulated with either ethanol alone or ethanol plus GSK3β inhibition. However, ethanol-induced regulation of PKA and presynaptic transmission networks was no longer present when animals were treated with tideglusib, suggesting GSK3β inhibition reverses transcriptional changes within these pathways.

An interesting protein which potentially connects both the PKA and presynaptic signaling networks is brain-derived neurotrophic growth factor (BDNF). BDNF has been shown to both reduce ADBE by inhibiting GSK3β[47] and activate PKA signaling, which also inhibits GSK3β[33, 67]. Chronic ethanol has been shown to decrease BDNF levels in the brain[68]. As such a potential pathway through which chronic ethanol decreases BDNF, resulting in less PKA-induced GSK3β phosphorylation and increased GSK3β activity might account for ethanol-induced neural adaptions, and these adaptations can be reversed through exogenous GSK3β inhibition.

The mechanism through which GSK3β induces these effects could be through Wnt signaling as evidenced not only through the results from our hierarchical clustering ontology analysis, but also from emergence of an expression network regulated by GSK3β inhibition in the presence of ethanol overrepresented for functions in transcription and multiple TFBS involved in both canonical and non-canonical Wnt signaling, including E12/TCF.

Additional studies expanding upon these results by looking at expressional networks within NAc could be particularly interesting. As stated, GSK3β inhibition was identified as reversing ethanol-induced mPFC expression changes in a network regulating presynaptic neurotransmission and excitatory/inhibitory balance. Given the known mPFC to NAc projections, we might expect to see more presynaptic alterations from axon terminals originating in mPFC projection neurons and potential postsynaptic transcriptional changes from dendritic synapses in NAc. Additionally, these analyses were limited to only male mice. Voluntary ethanol consumption is consistently higher in female animals, and we have recently demonstrated that tideglusib reduces ethanol consumption and preference in both sexes [15].

Furthermore, given the nature of these experiments, it cannot be known whether the ethanol-induced transcriptional changes observed are due to chronic ethanol itself or from abstinence in animals habituated to ethanol. Tissue samples for RNAseq were collected 24 hours after last ethanol access, meaning the animals were in a state of acute withdrawal.

Studies assessing transcriptional changes while ethanol is onboard could potentially elicit different results such as effects in pathways encoding for the rewarding properties of ethanol. However, assessing the state of the transcriptome during a withdrawal state may be more clinically relevant for AUD than during active intoxication. AUD develops progressively as the balance between the positive and negative reinforcing properties of alcohol shift[69]. Early alcohol exposure elicits positive reinforcement through lowering hedonic reward thresholds. With chronic exposure, homeostatic shifts occur over time to increase this threshold, producing decreased positive reward over time and eliciting aversive physical and affective symptoms during withdrawal, and this withdrawal has been shown to become progressively worse over time[69, 70]. As severity of withdrawal symptoms increases, the negative reinforcement provided by ethanol to relieve these symptoms also increases and is thought to be the main motivator for development of compulsive consumption and alcohol dependence[69]. As such, the ethanol-regulated gene networks identified here may encode for the motivation to consume alcohol during withdrawal states and GSK3β inhibition may reduce ethanol consumption through inhibition of these molecular adaptations.

There also remains the question of whether transcriptional differences observed between ethanol drinking vehicle-treated animals versus tideglusib-treated animals are due to direct effects of GSK3β activity on the expression networks or a result of the decreased ethanol consumption over time, resulting in a reduction in maladaptive ethanol-induced transcriptional changes. If network regulation was due to direct effects of GSK3β inhibition, we might expect to see networks regulated by tideglusib in the absence of ethanol, such as in our HT animals. However, there were no significant EW-dmGWAS networks in these ethanol naïve animals. As such it seems GSK3β inhibition may be regulating gene expression networks in ethanol consuming animals primarily through impacting ethanol-induced transcriptional changes that occur following repeated bouts of high ethanol consumption.

## ACKNOWLEDGEMENTS

Work was supported by grants from the National Institute for Alcohol Abuse and Alcoholism (NIAAA): R01AA027581 and P50AA022537 to MFM and F31AA030216 to SG

## Abbreviations

GSK3β: glycogen synthase kinase 3-beta
PFC: prefrontal cortex
AUD: alcohol use disorder
IEA: intermittent ethanol access
ANOVA: analysis of variance

